# Heritability and phenotypic plasticity of biting time behaviors in the major African malaria vector *Anopheles arabiensis*

**DOI:** 10.1101/2021.05.17.444456

**Authors:** Nicodem J. Govella, Paul C. D. Johnson, Gerry F. Killeen, Heather M. Ferguson

## Abstract

Use of Insecticide Treated Nets for malaria control has been associated with shifts in mosquito vector feeding behavior including earlier and outdoor biting on humans. The relative contribution of phenotypic plasticity and heritability to these behavioural shifts is unknown. Elucidation of the mechanisms behind these shifts is crucial for anticipating impacts on vector control. We used a novel portable semi-field system (PSFS) to experimentally measure heritability of biting time in the malaria vector *Anopheles arabiensis* in Tanzania. In PSFS assays, the biting time of F2 offspring (early: 18:00-21:00, mid: 22:00-04:00 or late: 05:00-07:00) was significantly associated with that of their wild-caught F0 grandmothers, corresponding to an estimated heritability of 0.30 (95% CI: 0.20, 0.41). F2 from early-biting F0 were more likely to bite early than F2 from mid or late late-biting F0. Similarly, the probability of biting late was higher in F2 derived from mid and late-biting F0 than from early-biting F0. Our results indicate that variation in biting time is attributable to additive genetic variation. Selection can therefore act efficiently on mosquito biting times, highlighting the need for control methods that target early and outdoor biting mosquitoes.

## Introduction

The most important current malaria control interventions are Insecticidal Treated Nets (ITNs) and indoor residual spraying (IRS) (1, 2). These tools have averted more than 600 million clinical cases in Africa since 2000 (1). The success of these interventions derives from their ability to exploit key aspects of the biting and resting behavior of African mosquito vectors, including their propensity to bite humans inside during sleeping hours, and rest inside after feeding (3, 4). The most important malaria vector species in Africa (*Anopheles gambiae* species complex (4, 5) and *An. funestus* (4, 5) typically exhibit these behaviours (6, 7). Despite the success of ITNs and IRS, their effectiveness is being undermined by mosquito responses that allow them to resist or evade them. Most notable is the widespread emergence of insecticide resistance (8). Additionally, there is growing evidence of shifts in vector behaviour in Africa and elsewhere (9–12), that allow them to reduce contact with ITNs and IRS (7). While the molecular and genetic basis of insecticide resistance (13, 14) and its impact on malaria transmission has been widely investigated (8, 15), much less is known about the basis of mosquito biting behaviour adaptations (16, 17) and their implications for vector control (3, 7).

Types of mosquito behavioural changes associated with ITNs and IRS include early-exiting from sprayed houses (18), increased outdoor biting at dawn or dusk when people are not protected by ITNs (10, 19–21) and increased feeding on livestock instead of people (22, 23). The capacity to mount such behavioural responses may vary between vector species. For example, historically *An. gambiae* in East Africa, has been reported to feed almost exclusively on people (24), inside houses and late at night (4, 6), while its sibling species *An. arabiensis* feeds more flexibly on humans and cattle (25, 26), indoors or outdoors, (27, 28), often in the early evening and dawn (27, 29). With the upscaling of ITNs, the relative abundance of *An. gambiae* compared to *An. arabiensis* has plummeted in several settings (30, 31). In west Africa, there are reports of a shift towards early evening or dawn biting, and more outdoor biting in *An. coluzzii* (32) and *An. funestus* (11, 19). Similarly, the proportion of “early” (18:00-21:00 hrs) and outdoor biting by the malaria vector *An. farauti* in the southwest Pacific increased after the implementation of IRS (10, 33). There is also evidence of shifts in host choice from humans to cattle in African malaria vectors following ITN and IRS implementation (22, 23). Shift in mosquito vector biting behavior directly impact the proportion of exposure that can be prevented by ITNs and IRS (3, 10).

Prediction of the impact of mosquito behavioural changes on vector control requires understanding of the mechanisms underlying them. It is unknown whether behavioural shifts reflect evolutionary adaptations in response to selection from ITNs/IRS, or are manifestations of pre-existing phenotypic plasticity. These possibilities are not mutually exclusive, but have different implications for control. The first hypothesis, defined as true *behavioral resistance*, is that behavioral traits are evolving in response to selection from interventions. Behavioral resistance traits could thus spread and become fixed in populations (3). The second hypothesis, defined as *behavioral resilience* (9), is that vector species were always capable of expressing alternative biting phenotypes, with this plasticity only exhibited in response to environmental variation that reduces human host availability. Here, biting behaviour may rapidly revert to baseline phenotype when control interventions are lifted (16). Behavioral resilience may define the limits of immediate behavioral responses to interventions (3), whereas behavioral resistance implies vectors can increasingly adapt their biting phenotypes to avoid indoor interventions over time; thus progressively eroding the proportion of exposure that can be prevented by their use. This mechanism may pose a larger problem than plasticity within a fixed range. For malaria elimination to be achieved, both phenomena will need to be tackled through adoption of complementary control methods that target vectors outside.

Here we made use of a portable semi-field system (PSFS) to experimentally investigate heritability and phenotypic plasticity in the biting time of the major African malaria vector *An. arabiensis* in Tanzania. These facilities enable mosquito host-seeking behavior to be measured under relatively realistic yet controlled and malaria-infection free conditions. Biting time phenotypes of wild-caught mothers (F0) were compared to phenotypes of their offspring (second generation) under controlled conditions.

## Methods

### Study site

All experiments were conducted in Lupiro village (−8.38 S, 36.67 E) within the Kilombero Valley, an area of moderate to high endemic malaria transmission in south-eastern Tanzania (34). Currently *An. arabiensis* is the most abundant malaria vector species in this area (35). Biting activity in this *An. arabiensis* population can start as early as dusk, with a peak around midnight and smaller peak toward morning (28). Most residents spend their time outdoors until ~10 pm when they go inside homes to sleep; with most using ITNs (28). All mosquito behavioural assays were conducted within a bespoke semi-field system, here referred to as a portable semi-field system (PSFS) installed temporarily in Lupiro Village (details in *Supplementary Information* **1**).

The PSFS was located in the same village where wild mosquitoes (parental generation) were collected to generate offspring for use in experiments.

### Assays for heritability in biting time

In Lupiro village, host-seeking female *An. arabiensis* were collected at different times of the night using Human Landing Catches (HLC). The collections were conducted in the peridomestic area around four houses (**SI 2**). In brief, volunteers collected mosquitoes hourly between 18:00–07:00hrs for two consecutive nights (14^th^ and 15^th^of July 2015). Mosquitoes collected in each hour were placed in separate paper cups. Mosquitoes visually identified as belonging to *An. gambiae sensu lato* (4, 5) from hourly collection were then grouped into one of 3 categories based on their time of capture: early (18:00-21:00), mid (22:00-04:00), and late (05:00-07:00) biting, and placed in separate holding cages. Biting activity was classified into these three discrete categories to correspond with times when people are likely to be either indoors and protected by ITNs (mid), or outdoors and unprotected (early and late). These mosquitoes formed the parental (F0) generation for experiments on biting time heritability. F0 females were provided with a blood meal for egg production as described in **SI 2**. Only eggs of mothers that were confirmed to be *An. arabiensis* by PCR were retained (**SI 2**). From these eggs, the F1 generation of each biting time phenotype category was reared separately to adulthood and used to produce an F2 generation for use in assays within the PSFS (**SI 2**). Experiments were conducted on the F2 rather than F1 generation to increase the number of offspring derived from each F0 biting time phenotype for experimentation.

Heritability was assessed by releasing F2 *An. arabiensis* into the PSFS, recording their biting time during a human landing catch and comparing it to that of their F0 grandmothers. One each night of experiment, 300 F2 *An. arabiensis* (100 from early, mid and late F0 phenotype) were released into the PSFS at 17:00 hours, with exception of one trial in which only 50 F2 from each three phenotypes were available (**SI 2**, and Statistical analysis methods below). Prior release, F2 mosquitoes were marked with either red, yellow or blue fluorescent dust colors according to their grandmothers’ biting time phenotype (18:00-21:00, 22:00-04:00, and 05:00-07:00). All marked mosquitoes were released simultaneously at the center within the PSFS. A volunteer entered the PSFS to conduct mosquito collections by HLC from 18:00 to 07:00 hrs (**SI 2**). Assays were conducted over 20 nights from 13^th^ August to 1^st^ October 2015 (**SI 2**). Additional HLC collections were conducted at local houses adjacent to the PSFS (within ~40 m) on the same nights as assays to confirm whether the temporal pattern of biting activity observed in the PSFS was consistent with that of the wild population. Wild mosquitoes were collected from inside and outside local houses.

### Data analysis

Before testing for heritability and association in biting time between F0 and F2, we first compared the nightly biting time profile of F2 offspring in the PSFS and that of wild *An. arabiensis* host seeking on the same nights as the experiments. This was to confirm whether the biting profile of mosquitoes inside the PSFS were representative of natural biting activity.

The proportion of mosquitoes caught biting each time period (early, mid and late) and its 95% CI were estimated separately in each location (indoor, outdoor and semi-field) by fitting logit-binomial GLMMs using the *glmer* function of the *lme4* R package (36), where biting time was modelled as a binomial response (early *vs* mid + late; mid *vs* early + late; late *vs* early + mid) and experimental night (20 nights of replicates) was fitted as an observation-level random effect. Bias in predicted proportions and 95% CIs due to Jensen’s inequality was corrected using the approximation of McCulloch et al (37).

Narrow sense heritability of biting time, *h^2^*, was estimated as *h^2^* = 2*t_F2-F0_*, where *t_F2-F0_* is the correlation between grand-offspring (F2) biting time and grandparental (F0) biting time (see **S 3 and S 4** for detailed methods and R code for estimation of *h^2^*. Owing to an unknown degree of assortative mating among the F1 generation, this *h^2^* estimate is expected to be positively biased, but we show that this bias is likely to be moderate to low (< 17% relative bias) for *h^2^* values below 0.5 (**S 3**). The correlation coefficient *t_F2-F0_* was estimated by modelling F2 biting time as an ordered categorical response (early < mid < late) in a mixed-effects ordinal probit regression model. This approach of modelling a discrete trait as the manifestation of an underlying continuous “liability” (here the tendency towards biting at a specific time) and estimating heritability on the liability scale is standard in quantitative genetics (38). F0 biting time was fitted as a fixed effect after being first converted to an integer score (early = 1; mid = 2; late = 3) then scaled to have unit variance and zero mean. Each experiment was conducted by releasing three (early, mid and late F2 biting) batches of mosquitoes on each of 20 days, motivating the inclusion of random effects for both date and batch within date to account for between-batch-within-day and day-to-day variation in F0 biting time. To allow for the potential effects of temperature and of the two volunteers who performed the HLC collections, these two factors were initially included in the model as fixed effects but were dropped because they showed no significant association with offspring biting time (P = 0.58 and P = 0.26 respectively).

In addition to estimating the heritability of biting time, we also tested whether individual F2 biting time phenotypes were associated with F0 biting time phenotype. Each F2 biting time phenotype was modelled as a binary response (early *vs* mid + late; mid *vs* early + late; late vs early + mid) in a logit-binomial GLMM using the *glmer* function of the *lme4* R package (36). F0 biting time was fitted as a categorical fixed effect, and the random effects fitted were date and an observation-level random effect. For each binomial F2 response the null hypothesis of no association with F0 biting time was tested using a likelihood ratio test. Pairwise differences in F2 biting time proportion between F0 biting times were tested using Wald tests. Predicted F2 biting time proportions with 95% confidence intervals (CI) were calculated from the fitted GLMMs. Bias in predicted proportions and 95% CIs due to Jensen’s inequality (39) was corrected using the approximation of McCulloch et al. (37).

### Ethical considerations

Prior to research commencing, approval for the study was obtained from the Institutional Review Board of Ifakara Health Institute in Tanzania (IHI/IRB/No. 16-2014), and Medical Research Coordination Committee of the National Institute of Medical Research in Tanzania (NIMR/HQ/R.8a/Vol.IX/1925). Meetings were held with local leaders in the village where the study was based, heads of households where mosquitoes were collected, and volunteers involved in collections, to explain the aims, advantages and potential risks of participating. Written informed consent was obtained from heads of households and volunteers participating in mosquito collections by HLC. All volunteers conducting HLCs were provided with malaria prophylaxis (Malarone^®^) during participation (40), and screened for malaria parasites by rapid diagnostic test) before and after participation. Participants involved in arm feeding of mosquitoes were first screened to ensure they were malaria parasite free and then received malaria prophylaxis (Malarone^®^) during experiments to protect them from infection and/or infecting mosquitoes (40). The study site was located close to a health clinic to facilitate rapid reporting and treatment of any malaria-related or other adverse events. No participants tested positive for malaria or reported any adverse effects from prophylaxis or any other procedure. Approval for publication was obtained from the Medical Research Coordination Committee of the National Institute of Medical Research in Tanzania.

## Results

Over 20 nights of sampling, 24,503 wild mosquitoes were collected, of which 28% (n=6883) were *An. gambiae s.l*. Of the 80% *An. gambiae s.l*. specimens that were successfully amplified by PCR, all were confirmed to be *An. arabiensis*. Of the 5850 F2 *An. arabiensis* released in the PSFS, 82% were recaptured ranging from 52% to 98% across experimental nights.

The biting time pattern of F2 *An. arabiensis* within the PSFS was similar to that observed in the wild population on the same nights (Fig. 1, Table. **SI 5**); confirming that representative biting behaviours were maintained in the PSFS. In all mosquito collections (made indoors, outdoors or in the PSFS), approximately one third of biting occurred in the early period of the night, with the bulk of activity occurring in the mid period and only a small proportion in the late period.

**Fig 1.**
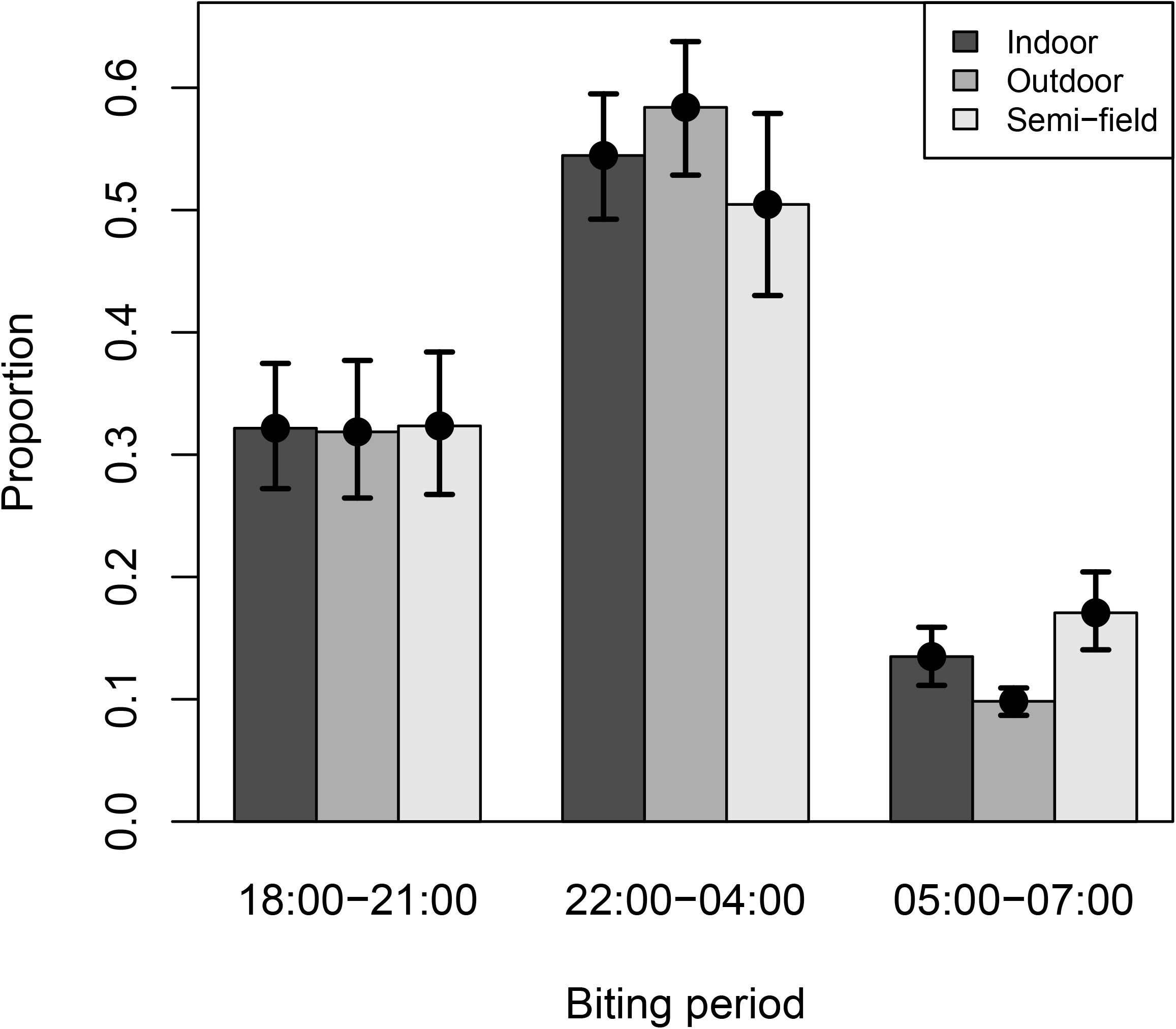
Proportion of *Anopheles arabiensis* biting at different periods of the night (early: 18:00-21:00; mid: 22:00-04:00, late: 05:00-07:00). Colors of the bars. Error bars are 95 CI from fitted model. .

### Heritability of biting time

A weak but highly significant positive correlation of 0.15 was estimated between the biting time phenotypes of F2 and F0 (*t_F2-F0_* [95% CI] = 0.15 [0.10, 0.21], p < 0.0001). Using the relationship *h^2^* = 2*t_F2-F0_*, this correlation translates to a biting time heritability of 0.30 [0.20, 0.41]. This heritability estimate is sufficiently low that the relative positive bias in the estimate due to assortative mating is likely to be less than 5% (**SI 3**), and therefore should not affect the conclusion that about a third of variation in biting time is due to additive genetic variation.

There was a significant association between F2 biting time, defined as a binary response, and F0 biting time in each of the binomial GLMMs (Table 1). The biting time of early and late biting F2 was positively associated with that of their F0 grandmothers (Fig. 2, Table 1). F2 from early-biting F0 were more likely to bite early than F2 of mid-biting F0 (P = 0.00061). F2 from mid-biting F0 were also more likely to bite early than F2 from late-biting F0 (P = 0.018) (Fig. 2, Table 1). A similar positive association was observed for the probability of biting late, with F2 from both mid (P = 0.029) and late-biting (P = 0.0014) F0 more likely to bite late than F2 from early-biting F0. However, offspring of mid-biting and late-biting F0 did not differ in their probability of biting late (P = 0.29). Most F2 biting activity occurred in the ‘mid period’ (Fig. 2). The probability of biting in the mid period did not differ between F2 from early and mid F0 (P = 0.066), and F2 from mid and late F0 (P = 0.19). However, F2 of late-biting F0 were more likely to bite in the mid period than those of early-biting F0 (P = 0.0017, Fig. 2, Table 1).

**Table 1.**
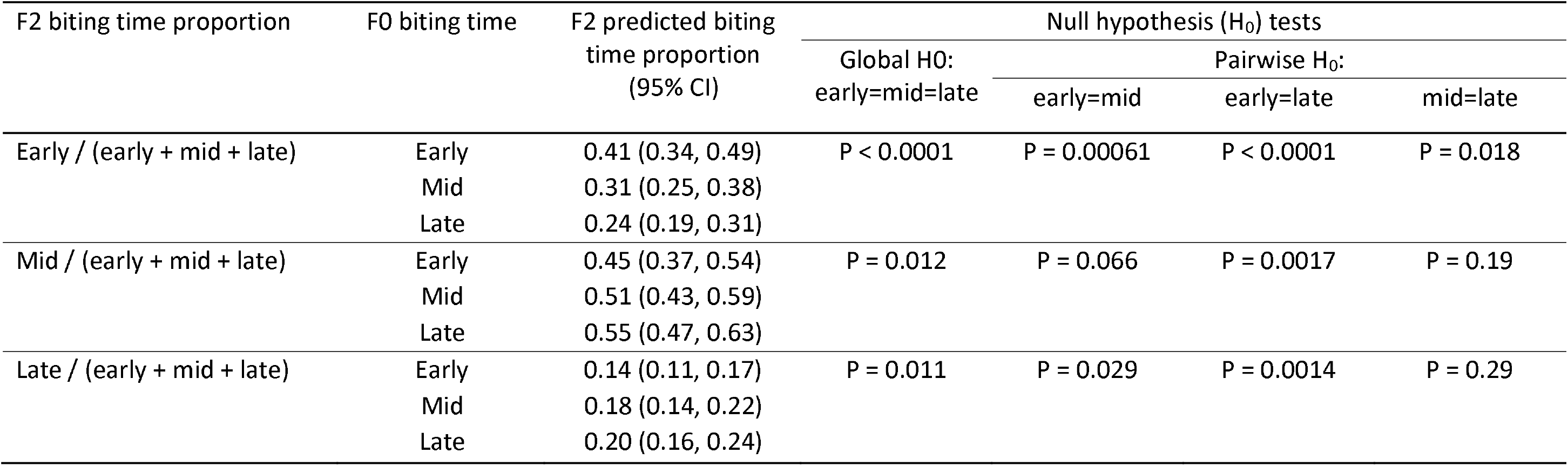
Proportions and 95% confidence intervals (95% CI) of F2 mosquitoes biting at early, mid and late periods of the night, estimated separately by the biting time of their F0 grandmothers. P-values for four null hypothesis tests (H0) are presented: the global null hypothesis that F2 biting time does not vary by F0 biting time, and the three pairwise null hypotheses of equal F2 biting time between early, mid and late F0 biting times. Biting time proportions and p-values were estimated using logit-binomial generalized linear mixed effects models (GLMM; see text for details).

| F2 biting time proportion | F0 biting time | F2 predicted biting time proportion (95% CI) | Null hypothesis (H<sub>0</sub>) tests |  |  |  |
| --- | --- | --- | --- | --- | --- | --- |
|  |  |  | Global H0:<br>early=mid=late | Pairwise H<sub>0</sub>: |  |  |
|  |  |  |  | early=mid | early=late | mid=late |
| Early / (early + mid + late) | Early | 0.41 (0.34, 0.49) | P < 0.0001 | P = 0.00061 | P < 0.0001 | P = 0.018 |
|  | Mid | 0.31 (0.25, 0.38) |  |  |  |  |
|  | Late | 0.24 (0.19, 0.31) |  |  |  |  |
| Mid / (early + mid + late) | Early | 0.45 (0.37, 0.54) | P = 0.012 | P = 0.066 | P = 0.0017 | P = 0.19 |
|  | Mid | 0.51 (0.43, 0.59) |  |  |  |  |
|  | Late | 0.55 (0.47, 0.63) |  |  |  |  |
| Late / (early + mid + late) | Early | 0.14 (0.11, 0.17) | P = 0.011 | P = 0.029 | P = 0.0014 | P = 0.29 |
|  | Mid | 0.18 (0.14, 0.22) |  |  |  |  |
|  | Late | 0.20 (0.16, 0.24) |  |  |  |  |

**Figure 2.**
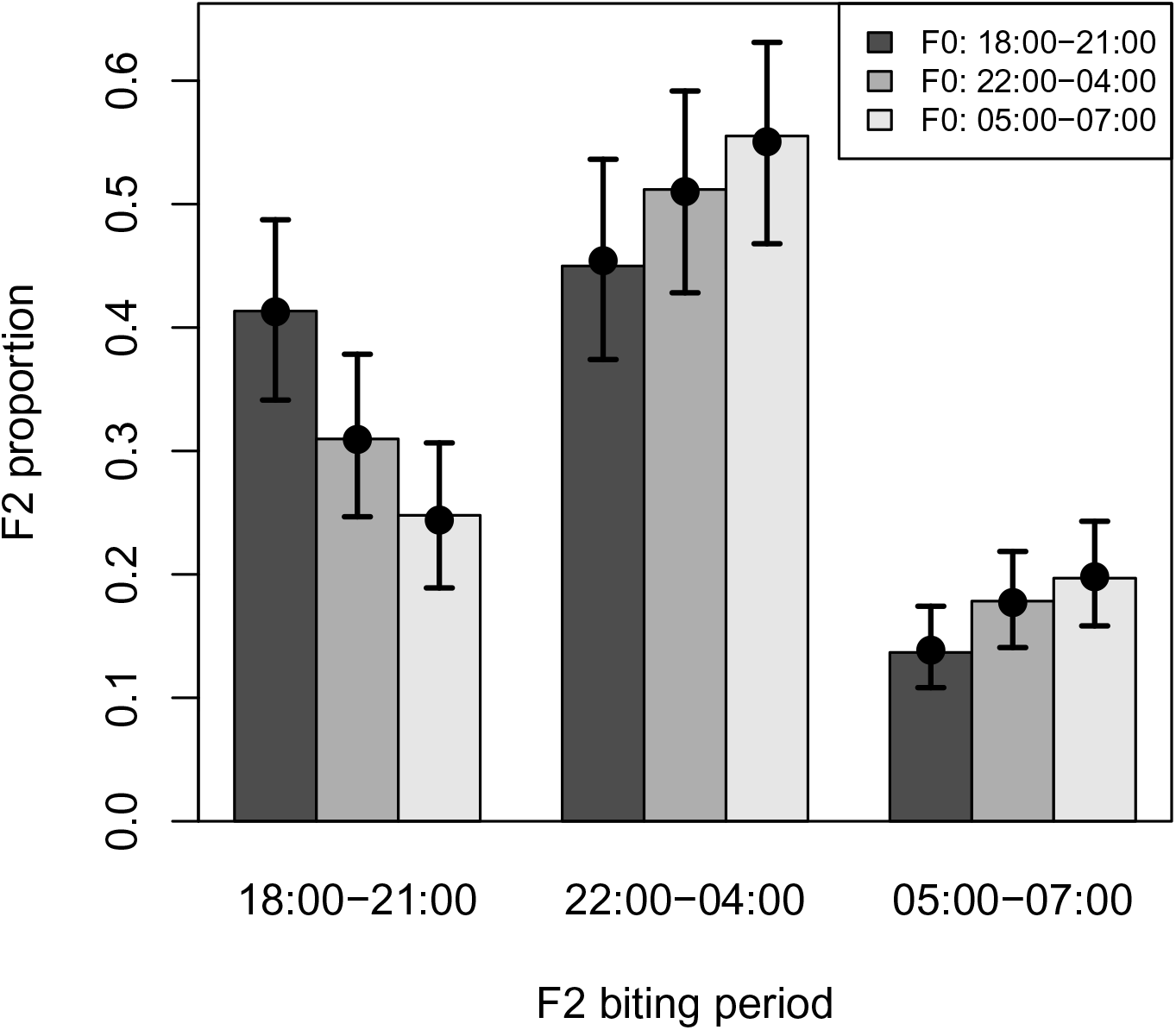
Predicted proportions of F2 biting at different times of the night relative to the biting time of their F0 grandmothers (early: 18:00-21:00hrs), mid: 22:00-04:00hrs, late: 05:00-07:00hrs). Colors of bars denote the biting time of F0 grandmothers. Error bars are 95% CI from the fitted model.

## Discussion

Here we experimentally investigated the heritable genetic basis of biting time in African malaria vectors through comparison of phenotypes in a wild F0 population and their F2 offspring, and found strong evidence of substantial genetic variation in biting times (41). Specifically, F2 offspring from early biting F0 grandmothers were more likely to bite early than offspring from mid or late biting F0. Similarly, F2 offspring from late biting F0 were more likely to bite late than offspring from early biting F0. In contrast the offspring of mid and late biting F0 were observed to bite in these two periods with similar frequency. Thus, the likelihood of an F2 feeding within either extreme of the biting activity range (either before 10pm or after 5am) was associated with their parental phenotype.

This evidence of a genetic association between “extreme” biting time phenotypes (e.g either early: 18:00-21:00 hours, or late 05:00-0.7:00) in offspring and grandparents indicates that selection generated by use of ITNs during typical sleeping hours (22:00-05:00hrs) could select for *An. arabiensis* to shift its biting time to periods when most people are unprotected. Here we deliberately chose to categorize mosquito biting times into periods of unequal length; early (4 hours), mid (6 hours) and late (3 hours), to focus analysis on heritability of behaviours that are specifically problematic for ITNs programmes. The extended length of the ‘mid’ period may account for why most *An. arabiensis* were observed biting during this time regardless of their grandparental biting time phenotype. Finer-scale shifts in mosquito biting time within the mid period may have little epidemiological consequence if people consistently use ITNs while sleeping; which typically occurs between 22:00-04:00) in African communities (28, 42). In contrast, a shift in mosquito biting to either before people go indoors to sleep or after they wake up in the morning would attenuate the impact of ITNs (7). For example, recent evidence from west Africa found *An. funestus* biting during morning hours (7am-11am) when people are outside in response to widespread use of ITNs (11, 19). Here we have shown that the tendency of *An. arabiensis* to bite during either of these extreme ranges does have a genetic component; raising concerns that malaria vectors may increasingly adapt their biting times under selection from ITNs. Such behavioural adaptability represents another effective mechanism of resistance against interventions in malaria vectors, beyond the usual physiological insecticides resistance traits, which may also contribute to both residual (7) and rebounding of transmission (3).

The timing of daily activities in other insects has been shown to have a genetic basis, including emergence in Drosophila (43), flight activity in Culex mosquitoes (44), locomoter activity in seed beetles (45) and mating in melon flies (46). Thus, circadian behaviours in insects are commonly under genetic control, with potential to respond to selection. There has been relatively limited investigation of the genetic basis of host seeking in mosquitoes; but one study found evidence of genetic structure between early and late biting *Nyssorhynchus (Anopheles) darlingi* in the western Amazon (47). In contrast, a previous investigation of genetic variation between early and late biting *An. arabiensis* in our study area found no evidence of substructuring within a range of candidate circadian genes and single nucleotide polymorphisms (48). Those authors acknowledged the study had low power to detect genetic variation for biting time due to limitations in sample size and the number of SNPs analyzed (48); and that a more robust association mapping analysis would be required to be conclusive. Our finding here of clear heritability in the biting times of *An. arabiensis* indicates that recently observed shifts in malaria vector biting time in response to ITNs (12, 49) may well result from extended evolutionary processes in addition to near-instantaneous phenotypic plasticity.

The study may have some limitations which could have prevented detection or biased results on heritability. An initial concern was that mosquito biting times observed under semi-field conditions may not be reflective of natural phenotypes. However, we confirmed that the nightly pattern of biting activity in the PSFS was similar to that observed in the local wild population on the same nights. Second, ability to detect heritability in biting time may have been reduced because instead of comparing mother-offspring pairs; analysis was based on pooled offspring generated from a cohort of F0biting in each time period. Thus, F2 biting time phenotypes represent an average phenotype from a group of grandmothers, and may mask stronger associations between specific parent-offspring pairs. Despite this limitation of the study design, trans-generational correlation in biting time were still unambiguously detected.

## Conclusions

This study provides the first experimental confirmation of heritability of biting time in *An. arabiensis*, one of the main malaria vectors in Africa. These results indicate that *An. arabiensis* can adapt its biting behaviour to avoid ITNs. Therefore, the observed changes in biting time by field populations may well be driven by intervention-induced longer-term selection. The role of genetics in this behavioural shift may be broader than captured here, and have direct implications for the maintenance of residual malaria transmission, and possibly rebounding of transmission in the future. This highlights the urgent need for complementary interventions that can target mosquitoes in outdoor environments, especially outside of typical sleeping hours.

## Acknowledgements

We thank the local leaders and residents of Lupiro Village for their cooperation throughout the study and to the volunteers for their commitment to this work. Special thanks go to Amos Thomas Mlwale, a technician for his tireless effort that ensured proper data collection from both field and semi-field experiments.

## Data availability statement

Access and use of data supporting this article will be made available in publicly repository.

## Author contributions

Conceptualization: NJG, GFK, HMF

Data curation: NJG

Formal analysis: NJG, PCJ

Funding acquisition: NJG

Investigation: NJG

Methodology: NJG, GFK, HMF

Project administration: NJG

Resources: NJG

Writing-original Draft preparation: NJG

Writing-Review & Editing: NJG, PCJ, GFK, HMF

## Authors declaration

All authors have read and approved the manuscript and that it has not been accepted or published elsewhere

## Supplementary information

This contains information and Figure related to Portable semi-field system (SI 1), Protocol and experimental design for assays of heritability in biting time (SI 2), Statistical methods for estimating the heritability of biting time (S3) and R-codes showing how assortative mating influence the phenotypic correlation among F1 mates and consequent expected bias in estimation of heritability (SI 4) and (SI 5) is raw-data summary comparing number caught from different biting periods and across different sources of collection.

