## Supplementary material for "Heritability and phenotypic plasticity of biting time behaviors in the major African malaria vector *Anopheles arabiensis*": SI 1. Portable semi field.docx

**SI 1: Portable Semi-field System (PSFS)**

A semi-field system, here referred as “Portable semi-field system” (PSFS), consisting portable netted cage ( 11.51 × 61.1 m with 1.85 m height was constructed (**Figs. SI** ). The floor of the PSFS was made of thick polyvinylchloride sheeting (PVC) which protects against rough substrate and surface water. The wall was composed of durable, Teflone-coated woven fiberglass netting to facilitate air movement and reduce inside temperature. The netting materials was chosen so that this system maintains natural weather conditions. The top of the netting cage is covered with canvas. This inner structure of the PSFS was also covered with an outer gabled roof made of collapsible pipes and supported with steel pipe posts distributed equidistant along the width and length. Along the gabled roof, there were sisal lopes running from all sides to the ground. This was made for tightening and increasing the stability of the PSFS against winds and heavy rains. Surrounding the floor, where this PSFS was placed in the field, a small concrete trench was established. This trench was always filled with water to stop ants from entering inside the PSFS, which would otherwise scavenging the mosquitoes. The entire process of installation or set up of the PSFS in the field may last within two to three hours. This PSFS was installed exactly at the same location where wild mosquito collections were conducted.

**Fig. SI 1. Floor plan and structure of the Portable, semi-field system.**

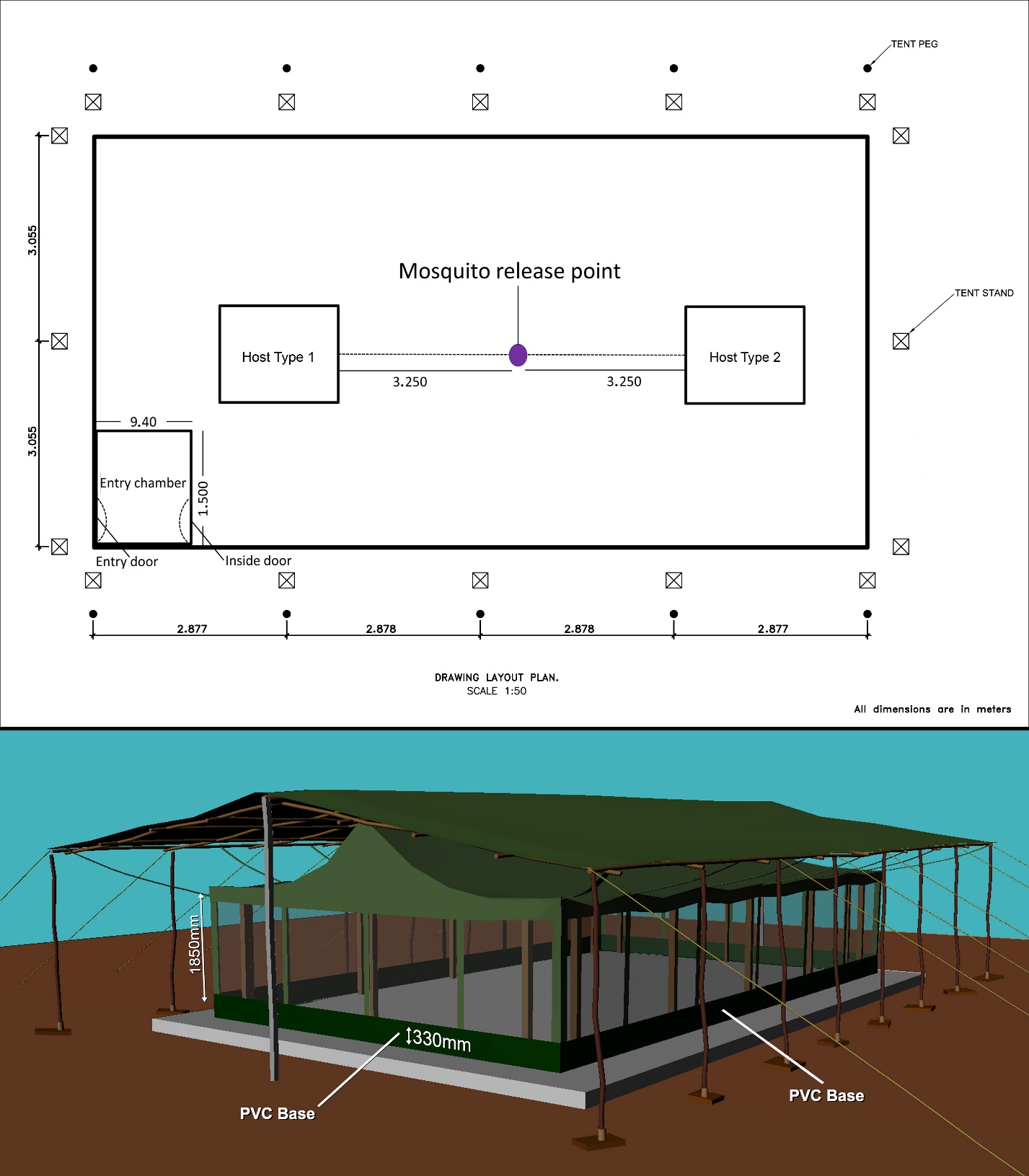
