## Supplementary material for "Heritability and phenotypic plasticity of biting time behaviors in the major African malaria vector *Anopheles arabiensis*": SI 2. Bioassays of biting time.docx

**Protocol and Experimental design for bioassays of heritability in biting time**

In Lupiro village, female *An. arabiensis* were collected host seeking at different times of the night using Human Landing Catches (HLC). The collections were conducted from peridomestic area at around 4 houses. With HLC, a volunteer exposes his lower limbs and aspirates mosquitoes landing on his exposed limbs. Volunteers collected mosquitoes host seeking between 18:00–07:00hrs for two consecutive nights (14^th^ and 15^th^ of July 2015) at 4 houses. Mosquitoes visually identified as belonging to *An. gambiae s.l.* (*Coetzee 2020 Malar J 19:70*) were grouped into one of 3 biting time periods on the basis of their collection time: early biting (18:00-21:00), mid biting (22:00-04:00), and late biting (05:00-07:00) and were placed in separate holding cages (0.36 m x 0.39 m). Biting time activity was broken into these 3 discrete categories to correspond with times when people are likely to be either indoors and protected by ITNs (‘mid’), or outdoors and unprotected (early and late). Cages were maintained under ambient conditions within the PSFS. Of the 245 female *An. gambiae s.l.* obtained, 218 survived the transition into cages (71, 98 and 49 from early, mid and late biting groups respectively). These mosquitoes formed the parental (F0) generation for experiments on biting time heritability. These mosquitoes were provided with a blood meal via arm feeding for two days consecutively post capture, with each group blood fed in the same time period they were collected. After blood feedings, members of the parental generation (F0) were individually placed into small, separate cages (0.15 m × 0.17 m) inside PSFS, and assigned a unique identification code (ID). Each small cage contained a petri dish lined with wet cotton and filter paper on top as an oviposition substrate. After oviposition, each individual mother was killed and stored on silica gel in a labeled 1.5 ml eppendorf tube. Females who produced eggs were killed and identified to species level by polymerase chain reaction (Scott et al 1993 *Am J Trop Med Hyg 49: 520-529*). The eggs from each confirmed *An. arabiensis* mother were transferred into a small water-filled bowl (~0.1 m diameter) for hatching and subsequent larval development. The larvae were fed on Tetramin fish food (Tetra, Melle, Germany). Pupae emerging from each egg clutch were combined into groups on the basis of the biting time category of their mother, and transferred into a larger holding cage for emergence (separate cages for each biting time phenotype). The first filial. generation (F1) adults emerging from these pupae were maintained on 10% glucose solution for up to 5 nights, and then given a blood meal via arm-feeding as described for the parental generation. This entire process was repeated to obtain a second generation (F2) for use in bioassays. Experiments were conducted on the F2 generation to increase the sample size in each biting time phenotype.

Heritability was tested using F2 *An. arabiensis* that were released in a PSFS and recaptured using HLC to measure their biting time phenotype, and compare it with that of their F0 grandmothers. Bioassays were conducted over 20 consecutive nights. One each night of bioassays, 300 F2 *An. arabiensis* (100 per biting time phenotype) were released into the PSFS at 17:00 hours, with the exception of one trial in which only 50 of each phenotype were available. Prior to release, F2 mosquitoes were starved for at least two hrs, and marked with either red, yellow or blue fluorescent dust colors according to their grandmothers’ biting time phenotype (18:00-21:00, 22:00-04:00, and 05:00-07:00). All marked mosquitoes were released simultaneously at the center within the PSFS. A volunteer entered the PSFS to conduct mosquito collections by HLC from 18:00 to 07:00 hrs, with mosquitoes attempting to feed during each hour grouped into a common holding cup. In the morning after collection, mosquitoes captured during each of the 3 biting periods were identified and recorded via by their dust color to their grandmother’s biting time. Thus both the biting time period of each individual could be linked to that of their grandmother. Additional HLC collections were conducted at 2 local houses adjacent to the PSFS within approximately 40m away on the same nights as bioassays to assess whether the pattern of biting activity in the PSFS was consistent with that of the wild population. Nearby houses from the PSFS were chosen so as to try maintaining similar environmental effects (for example winds, temperature and lights) between field and in the PSFS, which might affect the behavioural outcomes of the mosquitoes (*Kampango et al 2011 Med Vet Entomol 25:240-246*). Wild mosquitoes were collected from inside and outside local houses. In the morning after night collections, captured mosquitoes were killed using ethanol, sorted, identified to morphological level as *An. gambies s.l.* and their numbers recorded according to the time of their capture. All Individuals mosquitoes of *An. gambiae s.l.* (field collected mosquitoes) were stored in the eppendorf tube for sibling species identification by PCR (Scott et al 1993 *Am J Trop Med Hyg 49: 520-529*.)
