## Supplementary material for "Heritability and phenotypic plasticity of biting time behaviors in the major African malaria vector *Anopheles arabiensis*": SI 3. Analysis of HeritabilityMethods.docx

Supplementary methods for estimating the heritability of biting time

*Estimation of heritability of biting time*

Narrow sense heritability of biting time, *h^2^*, was estimated from the correlation between grand-offspring (F2) biting time and grandparental (F0) biting time, *t_F2-F0_*. This correlation coefficient was estimated using a mixed-effects ordinal probit regression model implemented using the *clmm* function in the *ordinal* package {Christensen, 2019 #5120} for *R* version 4.0.2. This approach of modelling a discrete trait as the manifestation of an underlying continuous “liability” (here the tendency towards biting at a specific time) and estimating heritability on the liability scale is standard in quantitative genetics {de Villemereuil, 2016 #5141}. F2 biting time, which is an ordered categorical response (early < mid < late), was modelled as resulting from trichotomizing a latent continuous biting time scale assumed to have a standard normal distribution (having zero mean and unit variance) around two threshold parameters represented by two intercepts in the regression model. F0 biting time was fitted as a fixed effect after being first converted to an integer score (early = 1; mid = 2; late = 3) then scaled to have unit variance and zero mean. Scaling both F2 and F0 biting times to unit variance makes the slope of the regression line equivalent to the F2-F0 correlation coefficient. A limitation of this model is that only F2 biting time was modelled on the liability scale, whereas the integer score of F0 biting time was effectively treated as an approximate direct measure of biting time liability. This process of approximation can be thought of as adding measurement error to the underlying F0 biting time, which would be expected to cause a moderate downwards bias in estimated heritability. Each experiment was conducted by releasing three batches (i.e early, mid and late biting F2) of mosquitoes on each of 20 days, motivating the inclusion of random effects for both date and batch within date to account for between-batch-within-day and day-to-day variation in F0 biting time, beyond the variation accounted for by the fixed effect of F2 biting time. To allow for the potential effects of temperature and of the two volunteers who performed the HLC collections, these two factors were initially included in the model as fixed effects but were dropped because they showed no significant association with offspring biting time (P = 0.58 and P = 0.26 respectively). Assuming random mating, *h^2^* = *t_F2-F0_*/*r*, where r is the coefficient of relatedness. The coefficient of relatedness between a single F0 grandmother and her F2 granddaughter is 0.25, giving *h^2^* = 4*t_F2-F0_*, but the F2-F0 correlation was contributed to by two grandmothers, giving *c*= 2*t_F2-F0_* {Falconer, 1996 #5122} However, this relationship applies only under random mating. Where mating is assortative, the phenotypic correlation between relatives should be accounted for, or the estimate of *h^2^* will be biased upwards {Nagylaki, 1978 #5123}. The degree of phenotypic correlation in this study is complex, and it’s potential impact on estimation of *h^2^* is discussed in the next section.

*Bias in estimated heritability due to assortative mating in the F1 generation*

In the F0 generation, grandmothers and grandfathers can be assumed to have mated at random. In the F1 generation, mating can be assumed to have been random, but within groups separated by grandmaternal biting time, which will have induced a phenotypic correlation that depends on *h^2^*. The effect of assortative mating is to increase the expected F2-F0 correlation so that in order to avoid a positive bias in estimation of *h^2^*, it should be divided by a factor of (1 + *r_P_*)(1 + *r_P_h^2^*), where *r_P_* is the phenotypic correlation between F1 mates {Nagylaki, 1978 #5123}. For example, for *r_P_* = 0.25 and a heritability of 0.7, a heritability that was estimated without accounting for assortative mating, as *h^2^* = 2*t_F2-F0_*, would be overestimated by 47%. We do not know the degree of assortative mating in this study, and therefore cannot adjust for it, but using simulations we show that *r_P_* is unlikely to be greater than 0.1 for *h^2^* < 0.5 (see below), in which case the positive bias in *h^2^* would be < 16%. We therefore estimated heritability using *h^2^* = 2*t_F2-F0_* but with the caveat that the low heritability estimates (< 0.5) are likely to be slightly positively biased and high (> 0.5) estimates could be severely positively biased due to deviation from random mating. In the next section we explore the likely extent of this bias in our biting time data.

*Estimation of bias in heritability due to assortative mating using simulated data*

We simulated biting time data of similar size and structure to the mosquito biting time data to explore how assortative mating influences the phenotypic correlation among F1 mates, and the consequent expected bias in estimation of heritability. 10,000 data sets were simulated across the full range of heritability values from zero to one. Grandparental biting time was simulated as an integer score, as in the data analysis. For computational speed, F1 biting time was simulated and modelled as a continuous directly measured scale rather than an ordinal manifestation of a liability scale. Also for computational speed, we avoided the need for random effects by simulating only a single pair of F1 offspring from each of 20 pairs of F0 grandmothers per biting time category (n = 60 F1 pairs). Assortative mating was simulated between F1 males and females by assuming random mating within the three groups defined by the F0 biting time of their grandmothers. To illustrate the effect of assortative mating, three outcome measures were plotted against true *h^2^*: the estimated phenotypic correlation (*r_P_*); the biased estimate of *h^2^*; and bias in estimating *h^2^* (Fig. 1-3). The smoothed relationship (i.e. averaging over sampling error in *r_P_*) between each outcome and true *h^2^* was estimated using OLS regression with a natural cubic spline with four degrees of freedom fitted to true *h^2^*. Full details of the simulation methods are given in the R code provided (**SI 4**).

The estimated phenotypic correlation among F1 mates was low (*r_P_* < 0.1) for *h^2^* < 0.5, rising to 0.5 as *h^2^* approached its maximum value of one (Fig. 1). Estimated *h^2^* including the expected bias due to not accounting for assortative mating was calculated as *h^2^*(1 + *r_P_*)(1 + *r_P_h^2^*) (Fig. 2), and percentage bias was calculated as 100 × ([biased *h^2^*] – [true *h^2^*])/[true *h^2^*] (Fig. 3). As expected, bias was low to moderate (0-17%) for *h^2^* < 0.5, but rose steeply (17-123%) for *h^2^* > 0.5. Estimated *h^2^* exceeded the maximum possible *h^2^* value of one for true *h^2^* > 0.7.

We conclude that estimates of biting time *h^2^* that are close to and above 0.5 could be moderately to severely positively biased by not accounting for the unknown degree of assortative mating, but that bias in estimates of *h^2^* below 0.5 is likely to be low (< 17%), and bias in *h^2^* estimates below 0.3 is likely to be negligible (< 4%).

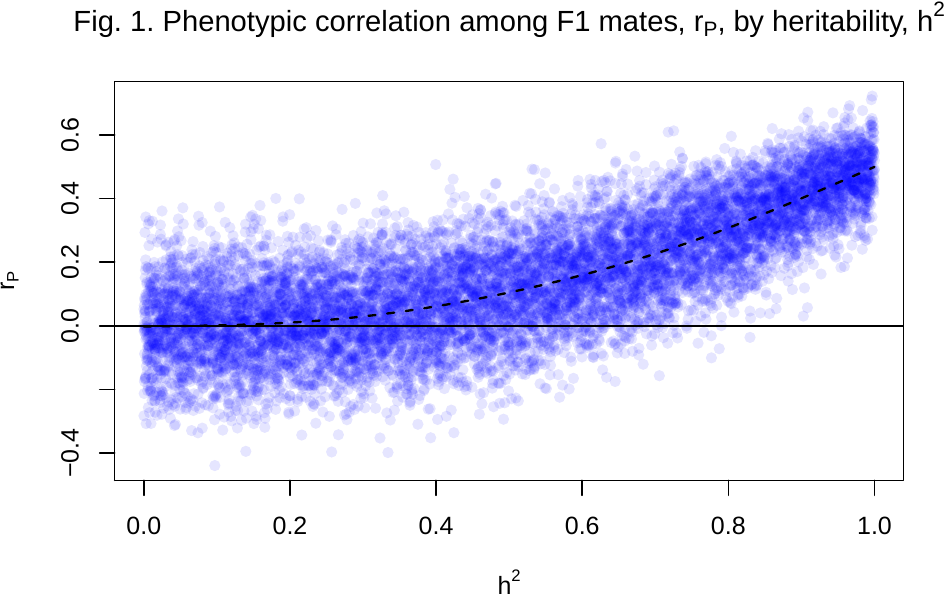

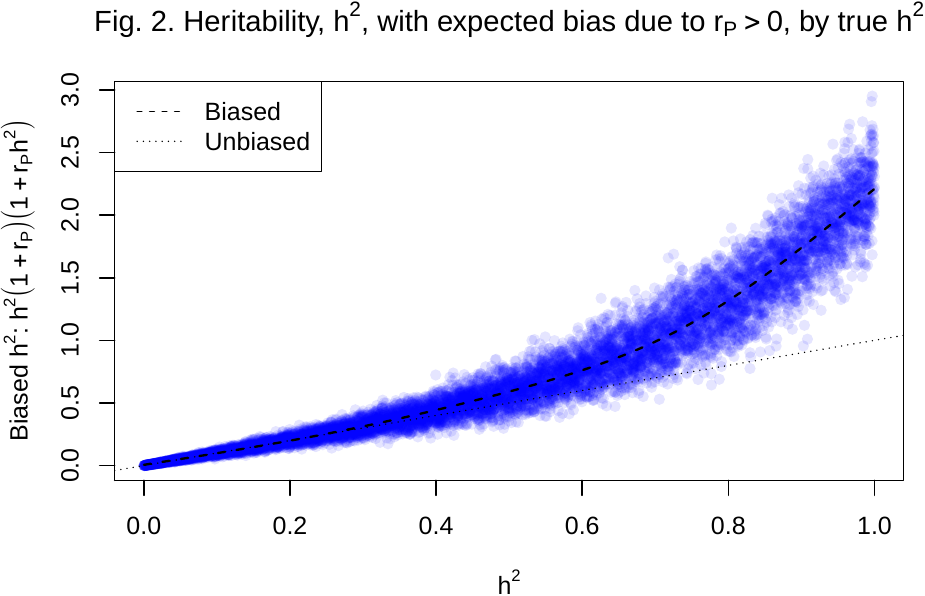

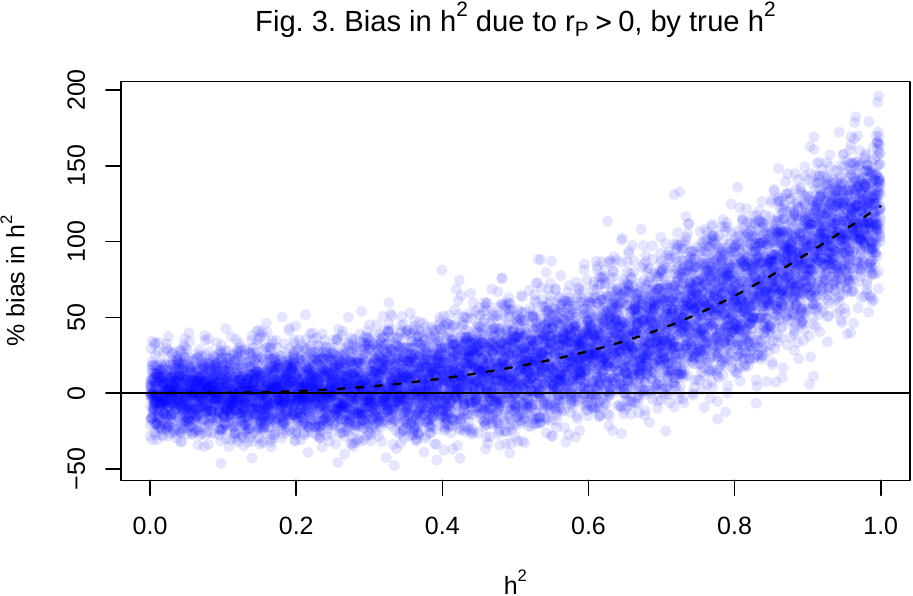
