## Supplementary material for "Heritability and phenotypic plasticity of biting time behaviors in the major African malaria vector *Anopheles arabiensis*": SI 4. Simulate.F1.mate.correlation.docx

Simulate.F1.mate.correlation.2020-11-22.R

#### pauljohnson 2020-11-23

**R script to support the calculation of heritability of mosquito biting time.**

*Background*

The aim of this script is to simulate data similar in size and structure to the mosquito biting time data, to show how assortative mating influences the phenotypic correlation among F1 mates, and the consequent expected bias in estimation of heritability.

*Start script*

Save figures to directory:

knitr**::**opts_chunk**$set**(fig.path = "Simulate.F1.mate.correlation.2020-11-22_figures/")

Load packages:

**library**(splines) **library**(scales)

Clear memory:

**rm**(list = **ls**())

*Simulation options*

Simulate assortative mating among F1 females and males? (change this to illustrate lack of bias when mating is not assortative).

assort <- TRUE

Number of offspring (similar to mosquito experiment):

n <- 60

Choose heritability values Simulate h2 values across its range from 0 to 1.

n.sim <- 10000

h2.list <- **seq**(0, 1 **-** 1**/**n.sim, length.out = n.sim) **+** 0.5**/**n.sim

Set random seed for repeatability [https://www.random.org/integers/?num=1&min=0&max=1000000000&](https://www.random.org/integers/?num=1&min=0&max=1000000000&col=1&base=10&format=html&rnd=new) [col=1&base=10&format=html&rnd=new](https://www.random.org/integers/?num=1&min=0&max=1000000000&col=1&base=10&format=html&rnd=new)

**set.seed**(567952879) *# Timestamp: 2020-11-23 11:13:18 UTC*

*Loop over h2 values estimating phenotypic correlation among F1 mates*

rP <-

**sapply**(h2.list, **function**(h2) {

*# variance parameters*

Vp <- 1 *# phenotypic variance*

Va <- Vp ***** h2 *# genetic variance*

Ve <- Vp **-** Va *# environmental/error variance*

*# start data frame to store values of relatives*

dat <- **data.frame**(id = 1**:**n)

*# simulate trait values (biting time) of grandparents (F0), and scale to standard normal* dat**$**GMm <- **as.vector**(**scale**(**sample**(**rep**(1**:**3, length.out = n)))) *# mothers of mothers* dat**$**GMf <- **as.vector**(**scale**(**sample**(**rep**(1**:**3, length.out = n)))) *# mothers of fathers* dat**$**GFm <- **as.vector**(**scale**(**sample**(**rep**(1**:**3, length.out = n)))) *# fathers of mothers* dat**$**GFf <- **as.vector**(**scale**(**sample**(**rep**(1**:**3, length.out = n)))) *# fathers of fathers*

*# make grandmothers perfectly correlated, as in the mosquito experiment # (this doesn’t affect the h2 estimate)*

**if**(assort) dat**$**GMf <- dat**$**GMm

*# simulate F1 values as the mean of their parents * h2 + environmental noise*

dat**$**F1m <- h2*****(dat**$**GMm **+** dat**$**GFm)**/**2 **+ rnorm**(n, sd = **sqrt**(Ve)) dat**$**F1f <- h2*****(dat**$**GMf **+** dat**$**GFf)**/**2 **+ rnorm**(n, sd = **sqrt**(Ve)) **var**(dat**$**F1m)

**var**(dat**$**F1f)

*# these lines can be run if stepping into the loop to show that # the simulated h2 values are accurately estimated:*

**if**(FALSE) {

*# estimate h2 by regression of F1 males on F0* fit <- **lm**(F1m **~ I**((GMm **+** GFm)**/**2), data = dat) **c**(h2 = h2, h2.est = **as.vector**(**coef**(fit)[2]))

*# estimate h2 by regression of F1 females on F0* fit <- **lm**(F1f **~ I**((GMf **+** GFf)**/**2), data = dat) **c**(h2 = h2, h2.est = **as.vector**(**coef**(fit)[2]))

*# ...these estimates can be shown to be accurate by choosing very large n, e.g. 10000*

}

*# permute F1 fathers to avoid correlations between F1 parents within F0 biting time*

*# groups (i.e. stratified by F0 grandmother’s phenotype). thus the only correlation is # due to stratification.*

dat**$**F1f.mix.i[dat**$**GMf **<** -1] <- **sample**(dat**$**id[dat**$**GMf **<** -1]) dat**$**F1f.mix.i[dat**$**GMf **>** 1] <- **sample**(dat**$**id[dat**$**GMf **>** 1]) dat**$**F1f.mix.i[**is.na**(dat**$**F1f.mix.i)] <- **sample**(dat**$**id[**is.na**(dat**$**F1f.mix.i)])

*# if no assortative mating then uncorrelate F1f from F1m*

**if**(**!**assort) dat**$**F1f.mix.i <- **sample**(dat**$**F1f.mix.i)

*# re-order F1f and GMf so that they are correctly aligned with offspring*

dat**$**F1f.mix <- dat**$**F1f[dat**$**F1f.mix.i] dat**$**GMf.mix <- dat**$**GMf[dat**$**F1f.mix.i]

*# correlation between F1 mates (rP)*

**cor**(dat**$**F1f.mix, dat**$**F1m)

})

*Plot results*

Function to plot a smooth spline (and optionally print a predicted value)

lines.spline <-

**function**(x, y, dof = 4, pred.x = NULL) {

**require**(splines)

fit <- **lm**(y **~ ns**(x, df = dof))

x.pred <- **seq**(**min**(x), **max**(x), length.out = 100)

**lines**(x.pred, **predict**(fit, **data.frame**(x = x.pred)), lty = 2, lwd = 1.5)

**if**(**!is.null**(pred.x)) {

**return**(**data.frame**(x = pred.x, y = **predict**(fit, **data.frame**(x=pred.x))))

}

}

Set up plotting parameters

col <- **alpha**("blue", 0.1) pch <- 16

Plot rP against h2 Add spline

### Fig. 1. Phenotypic correlation among F1 mates, r_P_, by heritability, h^2^

**plot**(h2.list, rP, xlab = **expression**(h**ˆ**2), ylab = **expression**(r[P]), col = col, pch = pch)

**lines.spline**(x = h2.list, y = rP)

**abline**(h = 0)

**title**(**expression**("Fig. 1. Phenotypic correlation among F1 mates, r"[P]*****", by heritability,

0.2

0.4

0.6

h"**ˆ**2))

r_P_

## 0.0 0.2 0.4 0.6 0.8 1.0

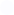

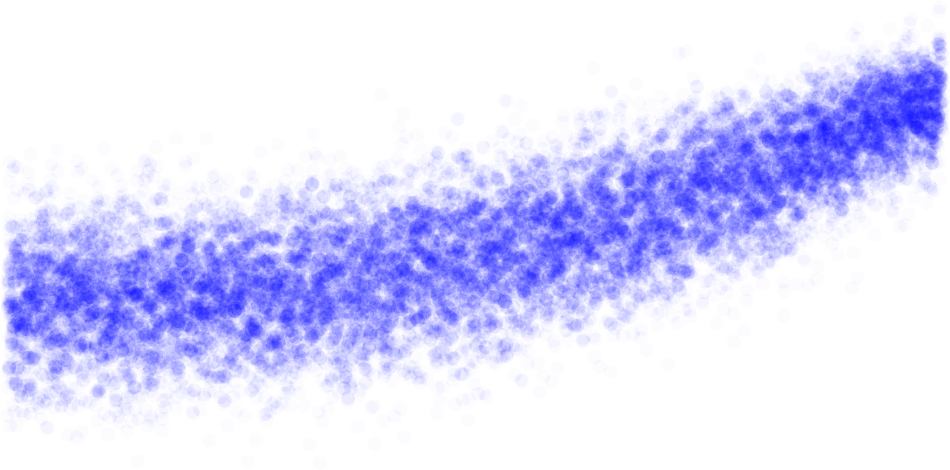

−0.4

0.0

h2

Plot h2 with expected bias against true h2 Add spline

h2.with.bias <- h2.list ***** (1 **+** rP) ***** (1 **+** rP ***** h2.list)

**plot**(h2.list, h2.with.bias, xlab = **expression**(h**ˆ**2), ylab = "",

col = col, pch = pch)

**mtext**(**expression**("Biased h"**ˆ**2*****": "*****h**ˆ**2 ***** (1 **+** r[P]) ***** (1 **+** r[P] ***** h**ˆ**2)), side = 2, line = 2.5)

1.5

**lines.spline**(h2.list, h2.with.bias)

**abline**(0, 1, lty = 3)

**title**(**expression**("Fig. 2. Heritability, h"**ˆ**2*****", with expected bias due to r"[P]**>**0*****", by true h"**ˆ**2))

**legend**("topleft", legend = **c**("Biased", "Unbiased"), lty = 2**:**3)

### Fig. 2. Heritability, h^2^, with expected bias due to r_P_ > 0, by true h^2^

2.0

2.5

3.0

## 0.0 0.2 0.4 0.6 0.8 1.0

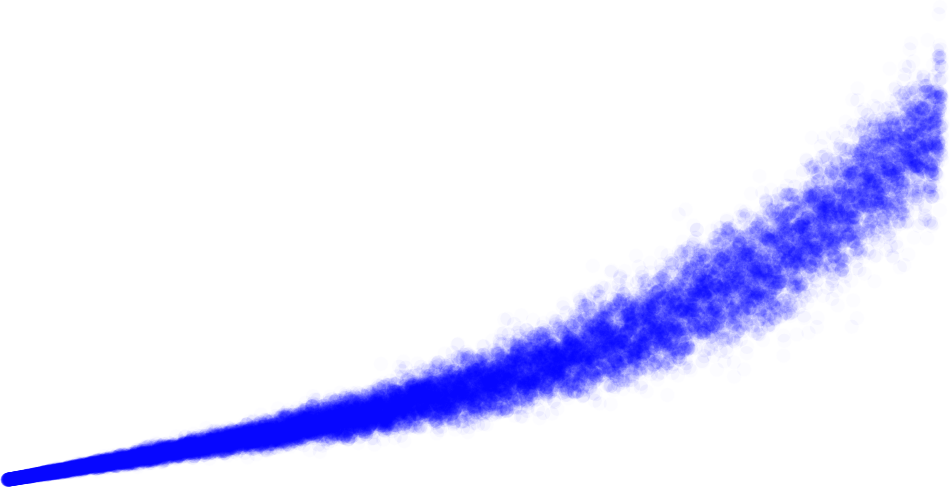

Biased

Unbiased

Biased h^2^: h^2^(1 + r_P_)(1 + r_P_h^2^)

0.0

0.5

1.0

h2

Plot percentage bias in h2 (given rP) against h2

Add spline and output expected bias in h2 at h2=0.5

h2.percent.bias <- 100 ***** (h2.with.bias**/**h2.list **-** 1)

**plot**(h2.list, h2.percent.bias, xlab = **expression**(h**ˆ**2), ylab = **expression**("% bias in h"**ˆ**2),

col = col, pch = pch)

**title**(**expression**("Fig. 3. Bias in h"**ˆ**2*****" due to r"[P]**>**0*****", by true h"**ˆ**2))

**abline**(h = 0) h2.percent.bias.at.h2 <-

**lines.spline**(h2.list, h2.percent.bias, pred.x = **seq**(0, 1, by = 0.1))

### Fig. 3. Bias in h^2^ due to r_P_ > 0, by true h^2^

100

150

200

## 0.0 0.2 0.4 0.6 0.8 1.0

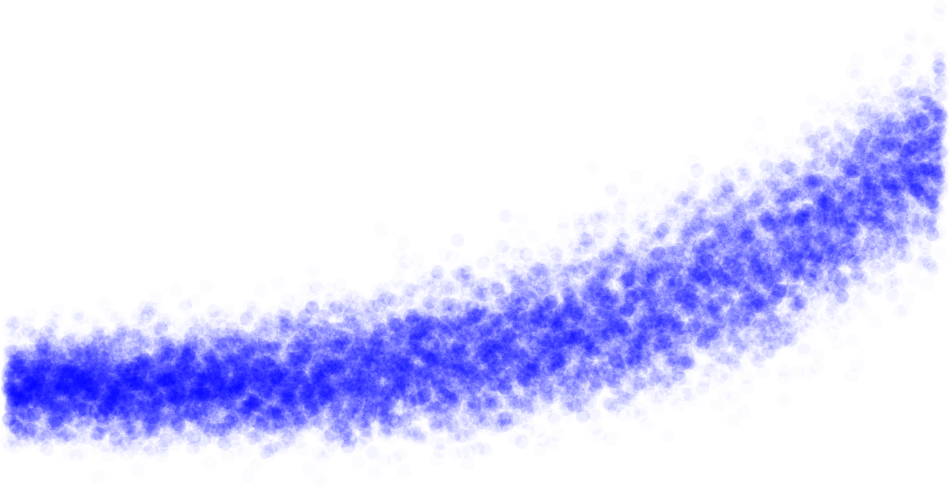

% bias in h^2^

−50

0

50

h2

**names**(h2.percent.bias.at.h2) <- **c**("h2", "percent.bias.in.h2.estimate") h2.percent.bias.at.h2**$**percent.bias.in.h2.estimate <-

**round**(h2.percent.bias.at.h2**$**percent.bias.in.h2.estimate)

Expected percentage bias in h2 when h2 = 0.3-0.5:

h2.percent.bias.at.h2	##	h2 percent.bias.in.h2.estimate

| ## | 1 | 0.0 | 0 |
| --- | --- | --- | --- |
| ## | 2 | 0.1 | 0 |
| ## | 3 | 0.2 | 1 |
| ## | 4 | 0.3 | 4 |
| ## | 5 | 0.4 | 10 |
| ## | 6 | 0.5 | 17 |
| ## | 7 | 0.6 | 28 |
| ## | 8 | 0.7 | 43 |
| ## | 9 | 0.8 | 64 |
| ## | 10 | 0.9 | 92 |
| ## | 11 | 1.0 | 123 |
