## Supplementary material for "Heritability and phenotypic plasticity of biting time behaviors in the major African malaria vector *Anopheles arabiensis*": SI 5. raw data comparison in biting time.docx

**Comparison in number caught from different biting periods and across different sources of collection**

| Biting time | Source of collection | Number caught | % Collected |
| --- | --- | --- | --- |
| 18:00-21:00 | PSFS | 1564 | 32.5 |
|  | Indoor | 576 | 31.0 |
|  | Outdoor | 1183 | 31.5 |
| 22:00-04:00 | PSFS | 2428 | 50.4 |
|  | Indoor | 1046 | 56.0 |
|  | Outdoor | 2200 | 59.0 |
| 05:00-07:00 | PSFS | 821 | 17.1 |
|  | Indoor | 252 | 13.4 |
|  | Outdoor | 370 | 10.0 |
